# Environmental impacts on gene expression noise and its relationship with fitness

**DOI:** 10.64898/2026.05.18.725919

**Authors:** Taslima Haque, Mohammad A. Siddiq, Fabien Duveau, Patricia J. Wittkopp

## Abstract

Genetically identical cells grown in the same environment show variation in gene expression known as expression noise. Expression noise can be heritable and impact fitness, making it subject to natural selection. Increasing expression noise for the *Saccharomyces cerevisiae TDH3* gene was shown to be beneficial in glucose-based media when mean *TDH3* expression was far from the fitness optimum but deleterious when it was close to this optimum. Here, we show that growth on different carbon sources alters the effects of new mutations on *TDH3* expression noise and examine the fitness effects of changing expression noise. In galactose-based media, we observed the same relationship between expression noise and fitness seen in glucose-based media, but in glycerol- and ethanol-based media, we observed the opposite relationship or no significant relationship, respectively. Using simulations of single-cell organisms, we found that these differences were most likely explained by environment-specific relationships between gene expression and fitness. We also found that, far from the optimum, the fitness effects of noise were greatest when expression was highly heritable between mother and daughter cells. The empirical observations and simulations reported in this study show how environments influence both the production of expression noise and its impacts on fitness.

## Introduction

An organism’s phenotype is determined by the interaction of genetic, environmental, and stochastic factors. The fitness of a genotype reflects how well its phenotype allows individuals to survive and reproduce in all the environments they experience. Phenotypic plasticity, which is itself a heritable and evolvable trait, allows the same genotype to produce different phenotypes in different environments [1]. Genotypes exhibiting phenotypic plasticity are selectively favored when this environment-specific phenotype increases survival and reproduction. But phenotypic plasticity is not the only adaptive strategy in dynamically changing environments; in such conditions, genotypes with higher phenotypic noise may also be favored by selection. [2]. This noise refers to the phenotypic variance among genetically identical individuals exposed to the same environment [3]. A genotype with higher noise may have a higher probability that, by chance, an individual with that genotype will be well-suited to a new environment, a strategy known as bet-hedging [2]. Because the extent of noise is also heritable and can affect fitness, it is also an evolvable trait [4,5]. Understanding the complex relationships among genetic variation, phenotypic plasticity, noise, and fitness in changing environments remains a pressing challenge for understanding and predicting biological evolution.

Phenotypic plasticity and noise at the organismal level can be generated at the molecular level by gene regulatory networks controlling gene expression [6]. Phenotypic plasticity can occur when a cell senses a change in environment and responds by altering gene expression [7], whereas phenotypic noise can result from variation in gene expression caused by the inherent stochasticity of molecular, chemical, and physical processes involved in the regulation of gene expression [4,8,9]. *cis*-regulatory sequences (e.g., promoters, enhancers) interacting with *trans*-regulatory factors (e.g., transcription factors) control when, where, and how much a gene is expressed. Genetic variation within both *cis*- and *trans*-regulatory components can alter mean expression levels, expression plasticity, and expression noise [5,7,10–12].

The role of *cis*-regulatory variation in shaping expression plasticity and noise has perhaps been studied most extensively in the baker’s yeast, *Saccharomyces cerevisiae*, where changes in the environment often cause widespread changes in gene expression [13]. This work has shown that *cis*-regulatory divergence affects not only a genotype’s average expression level but also its plasticity and expression noise (e.g., [12,14,15]). In some cases, the architecture of the promoter (which in *S. cerevisiae* includes both the basal promoter and upstream regulatory sequences) can cause mutations to have correlated effects on expression plasticity and noise [16]. Differences in expression noise among genes have been shown to correlate with promoter properties [9,11,17–19], the fitness effects of changing expression of a gene [20], and growth rates [21]. Understanding how these genome-wide patterns of expression noise translate into specific fitness consequences for individual genes requires a comprehensive and integrated analysis of the effects of genetic variants on expression level, expression noise, plasticity, and fitness in multiple environments.

The *S. cerevisiae TDH3* gene, which encodes a glyceraldehyde-3-phosphate dehydrogenase (GAPDH) protein that functions in glycolysis and gluconeogenesis, has been used extensively to study the effects of genetic variants on gene expression [22]. Previous work has characterized the effects of genetic variants in the *TDH3* promoter on expression level [5,23], expression noise [5,24], expression plasticity [10,25], and fitness [24,26], making it one of the most thoroughly characterized promoters in terms of the relationships among genotypes, phenotypes, and fitness in any organism. Studies comparing the effects of mutations in the *TDH3* promoter among environments have used media with different primary carbon sources (i.e., glucose, galactose, glycerol, ethanol). Cells growing by fermentation on glucose or galactose produce ethanol and glycerol as byproducts that can subsequently be used as carbon sources for respiratory growth [27]. Consequently, these environments provide biologically relevant contexts for examining how the environment shapes expression noise and its fitness consequences [28]. This work has shown that *TDH3* expression is plastic, with expression levels differing among these environments [10], and that the relationship between *TDH3* expression and fitness is also environment-specific [25]. However, the way these mutations affect expression noise in other environments (i.e., the plasticity of expression noise), as well as the way differences in the relationship between *TDH3* expression level and fitness among environments affect the fitness effects of changing expression noise, remain unknown.

To address these knowledge gaps, we used the data described in Duveau *et al*. [24] and Siddiq *et al. [25]* to measure how expression noise for 47 different *TDH3* promoter alleles differs for cells grown in media with glucose, galactose, glycerol, or ethanol as primary carbon sources. We then determined how the variation in expression noise introduced by these promoter alleles altered fitness in each environment independently from their effects on average expression level. These analyses showed that contexts in which increasing expression noise was beneficial differed between fermentable and non-fermentable environments. Simulations suggest that these differences are best explained by differences in the shape of the fitness function relating *TDH3* expression levels to fitness (i.e., doubling times for a single-celled organism) among environments. Potential differences in heritability of expression levels between mother and daughter cells, which could vary among environments, were also found to impact the relative fitness of different genotypes. However, differences in heritability were unable to explain why expression noise was beneficial at different average expression levels in different environments. Rather, the fitness effects of increasing expression noise were best explained by the concavity of the local fitness function, consistent with Jensen’s inequality from mathematics [29]. Taken together, this work demonstrates that the fitness effects of expression noise are environment-dependent in ways that are governed by the shape of the relationship between gene expression level and fitness.

## Results and Discussion

### Plasticity in expression noise driven by the *TDH3* promoter varies among environments

To understand how genetic variation in the *TDH3* promoter affects expression noise in different environments, we used data from 47 strains of *S. cerevisiae*, each containing a different allele of the *TDH3* promoter (*P*_*TDH3*_) driving expression of a yellow fluorescent protein (YFP) integrated into the genome at the *HO* locus [24,25]. These 47 strains include 39 with one or more mutations in known functional elements of the *TDH3* promoter (i.e., RAP1 or GCR1 binding sites or the TATA box, Figure 1A) in the *P*_*TDH3*_*-YFP* reporter gene, as well as 8 strains with two copies of a *P*_*TDH3*_*-YFP* reporter gene containing similar mutations (Supplementary Table 1). Data from 4 control strains was also used in our analysis (separate references were used for strains with one or two copies of the *P*_*TDH3*_*-YFP* reporter; Supplementary Table 1). In each of these strains, the native *TDH3* gene was unmutated (Figure 1B). Expression of YFP, as a proxy for activity of the *TDH3* promoter allele(s) in the reporter genes, was measured for at least 20,000 individual cells per strain in 3-4 replicates in each of 4 different environments using flow cytometry (Figure 1C). These four environments were media containing glucose, galactose, glycerol, or ethanol as the carbon source (see Methods). After correcting for differences in cell size, the fluorescence values from each replicate provide a distribution of expression levels for single cells in that replicate. For each strain in each environment, the median and standard deviation were calculated for each replicate (Figure 1D; Supplementary Table 2-5) and averaged among replicates (Supplementary Tables 6).

**Figure 1:**
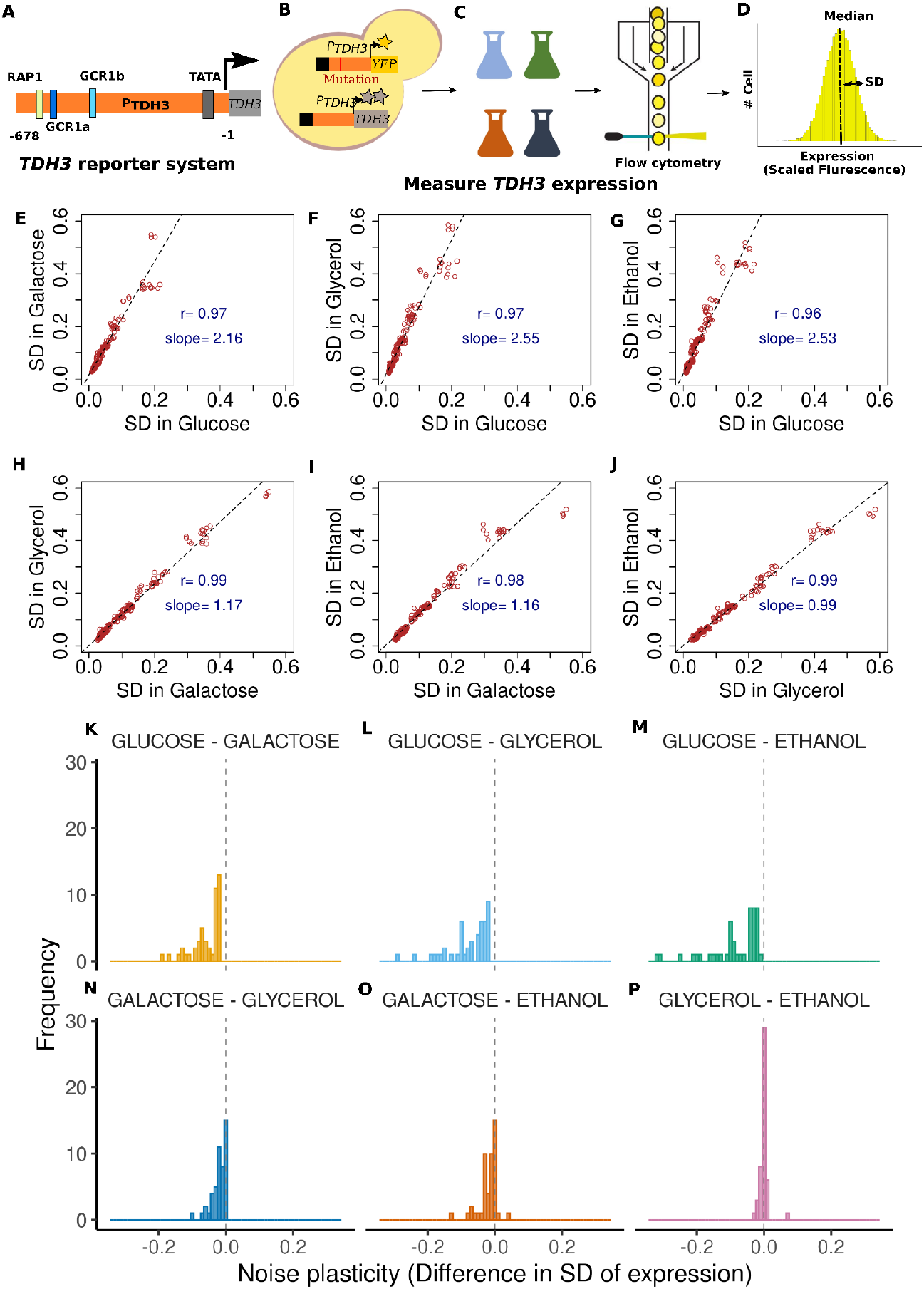
Plasticity in expression noise across different environments: **A)** Schematic diagram of the *TDH3* promoter in *S. cerevisiae* showing locations of binding sites for the RAP1p and GCR1p transcription factors and the location of the TATA box (adapted from Siddiq *et al*. [25]). **B)** An illustration of mutant strains used to measure YFP fluorescence driven by the *TDH3* promoter as a proxy for *TDH3* expression. Each mutant strain has a *P*_*TDH3*_*-YFP* reporter gene at the *HO* locus carrying a mutation(s) in the promoter and a wild-type *P*_*TDH3*_ at the native *TDH3* locus. C) The four colored flasks represent the four different types of media, two fermentable (glucose- or galactose-based) and two non-fermentable (glycerol- or ethanol-based), used in this study. A flow cytometer schematic is also shown measuring fluorescence for individual cells. D) Distribution of expression levels for 20,000 cells collected from an example strain, with median expression level and standard deviation (expression noise) labeled (adapted from Duveau *et al*., [24].). **E-J)** Expression noise driven by the *TDH3* promoter is strongly correlated between all six pairs of the four environments tested. The standard deviation of YFP expression is plotted as a measure of expression noise for the 47 *P*_*TDH3*_*-YFP* reporter genes in each pairwise comparison. The X and Y axes indicate the specific environments (glucose-, galactose-, glycerol-, or ethanol-based media) compared in each plot. Dotted lines show the linear regression for each pair of environments, with the regression coefficient and slope indicated on each plot. **K-P)** The distributions show pairwise differences in noise for the six different environmental contrasts.

Using standard deviation to measure expression noise for the *P*_*TDH3*_*-YFP* reporter gene, we found that expression noise differed among the 47 *TDH3* promoter alleles in each of the four environments (glucose: ANOVA, *F =* 2355.9, *df* = 47, *P =* 2.5 × 10^-219^; galactose: ANOVA, *F =* 2532.3, *df* = 47, *P* = 5.14 × 10^-222^; glycerol: ANOVA, *F =* 1058.8, *df* = 47, *P =* 1.52 × 10^-189^; and ethanol: ANOVA, *F =* 2355.9, *df* = 47, *P =* 2.53 × 10^-219^). This observation confirms that genetic variation alters not only the average expression level but also expression noise in each of the four environments. A significant, positive correlation was seen between the effects of mutations on average expression level and expression noise in all four environments (Pearson’s correlation coefficient > 0.88, p-value < 2.22 × 10^-16^), as expected from prior work on mutations in the *TDH3* promoter [23] and in other *S. cerevisiae* promoters [9,11]. Although we focus on standard deviation as a measure of expression noise throughout the main text, multiple metrics have been used to describe expression noise, and we found that the differences in expression noise in each environment were also significant when the standard deviation was scaled by the median expression level to produce a value similar to the coefficient of variation (CV) and when the standard deviation was squared (i.e., to calculate the variance) and then scaled by the median expression level to produce a value similar to the Fano factor (F) (P < 0.05; Supplementary Table 7).

The effects of the 47 different *TDH3* promoter alleles on expression noise were highly correlated in all six pairwise comparisons of the four environments (Pearson’s *r* > 0.96 in all cases; Figure 1E-J). However, a linear model for expression noise with strain and environment as factors showed a significant interaction term (F-value = 181.36, df = 141, p-value < 2.22 × 10^-16^), and the slope was significantly different from 1 in five of the six pairwise linear regression analyses (two-sided t-test p-value < 0.0083; Figure 1E-J). These findings show that the environment often influences the expression noise of a genotype. The same pattern was seen using other measures of expression noise (Supplementary Figures 1 and 2). Consistent with these observations, comparing expression noise for each of the 47 *P*_*TDH3*_*-YFP* mutant strains and the unmutated reference strain between pairs of environments often showed statistically significant differences in expression noise based on *t*-tests with a Bonferroni-corrected significance threshold of p < 0.00017 (Supplementary Table 8).

Expression noise tended to be lower in glucose-based media than in any of the other three types of media (Figure 1K-M; one-sample t-tests, H_0_ = 0, p_glu-gal_ = 6.84 × 10^-41^, p_glu-gly_ = 4.67 × 10^-41^, and p_glu-eth_ = 4.26 × 10^-41^). Indeed, 100%, 100%, and 96% of the 48 strains showed significantly lower expression noise for cells grown in glucose-based media compared to cells grown in galactose-, glycerol-, or ethanol-based media, respectively (Supplementary Table 8). Because cell populations grew faster in glucose-based media than in any of the other three environments tested (doublings per hour reported in [25]) as 0.72 in glucose, 0.56 in galactose, 0.36 in glycerol, and 0.25 in ethanol), these data suggest that expression noise tends to be lower when cells divide faster. Consistent with this idea, we also found that expression noise tended to be lower in galactose than in glycerol or ethanol (one-sample t-tests, p_gal-gly_ = 2.50 × 10^-30,^ p_gal-eth_ = 2.52 × 10^-22^; Figures 1N and O); however, these differences in expression noise tended to be smaller than in the comparisons to glucose. Only 48% and 44% of the strains showed statistically significant differences in expression noise when comparing cells grown in galactose with cells grown in glycerol or ethanol, respectively (Supplementary Table 8). For cells grown in glycerol- and ethanol-based media, we saw no systematic difference in expression noise (one-sample t-test, p_gly-eth_ = 1, Figure 1P) despite a difference in growth rate, and only 13% of strains showed a significant difference in expression noise (Supplementary Table 8).

### Fitness effects of increasing *TDH3* expression noise vary among environments

To examine the fitness effects attributable to expression noise in different environments for *TDH3*, we used data from a different set of 47 strains (plus relevant control strains) that are similar to those described above for expression analysis except that the mutations were introduced at the native *TDH3* locus instead of in the *P*_*TDH3*_*-YFP* reporter gene (Supplementary Table 9, Figure 2A). The relative fitness of strains with mutations at the native *TDH3* locus was determined from head-to-head competitions between each mutant strain (marked with YFP) and a common, unmutated reference strain (marked with GFP) (Figure 2A-C; Supplementary Table 10). These measures of relative fitness were then combined with the expression data described above to generate environment-specific fitness functions that describe how changes in average *TDH3* expression level impact fitness. As originally shown in Siddiq *et al. [25]* and recreated in Figure 2D, the relationship between average *TDH3* expression level and fitness differs in each of the environments. The fitness functions in the two fermentable carbon sources (glucose and galactose) had similar shapes, but the optimal expression level for galactose was lower than for glucose. The fitness function for glycerol displayed a wider plateau around the optimal expression level than for glucose or galactose, with a slightly steeper decrease of fitness for expression levels outside this plateau. The fitness function in ethanol was the flattest, indicating that this environment showed the smallest differences in fitness among cells with different levels of *TDH3* expression. It was also the most rugged, with a local optimum (Figure 2D).

**Figure 2:**
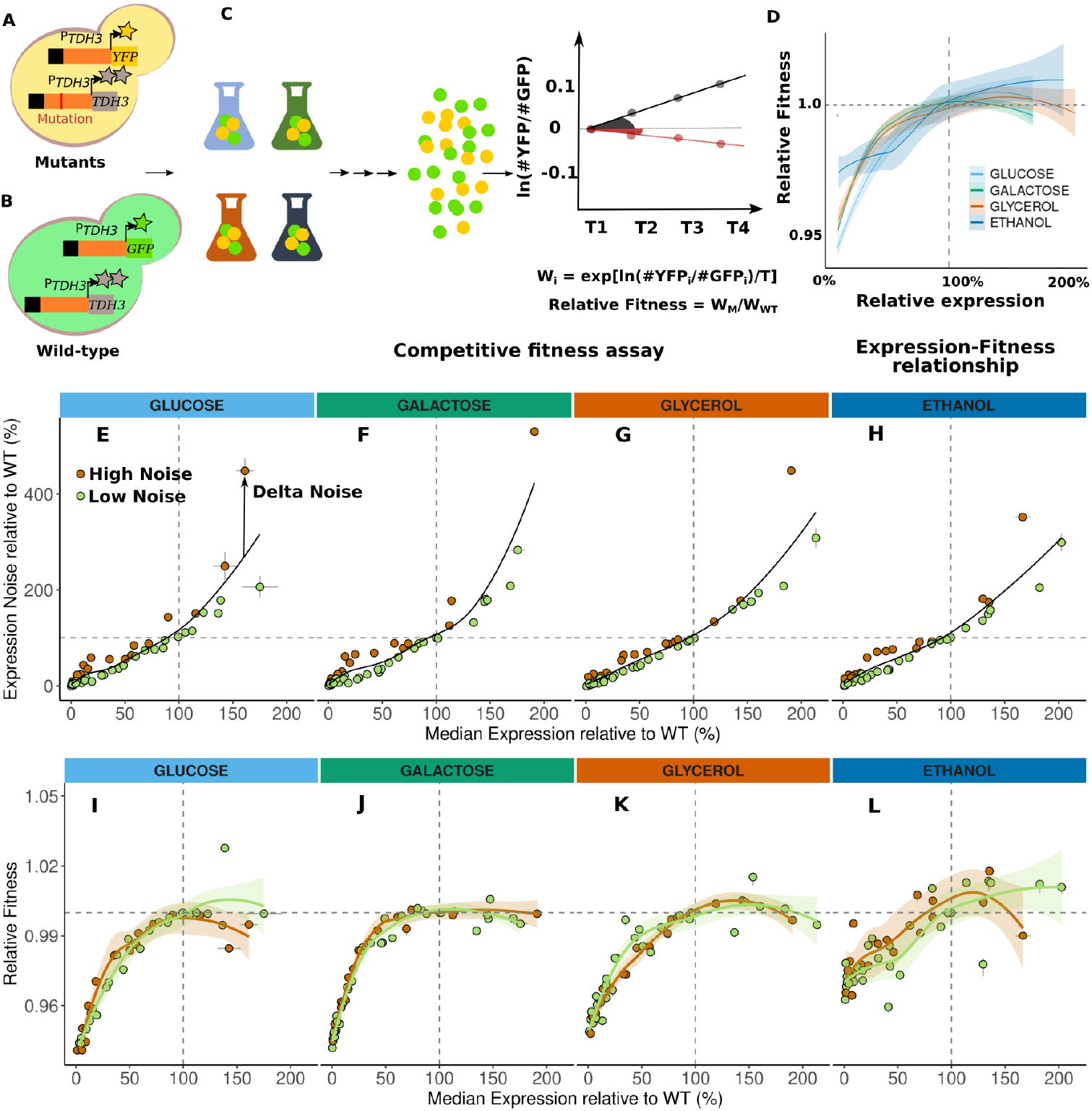
Relating relative fitness to expression level of *TDH3*. A-C) Schematics show the competitive growth assay used to measure the relative fitness of *TDH3* promoter mutant alleles (adapted from Duveau *et al*. [24]). Each mutant genotype contains a *P*_*TDH3*_*-YFP* reporter gene at the *HO* locus and a single *TDH3* promoter mutation at the native locus (A). The common competitor genotype has a *P*_*TDH3*_*-GFP* reporter gene at the *HO* locus and a wild-type *TDH3* promoter at the native locus (B). Competitive growth assays were carried out in media containing glucose (light blue), galactose (green), glycerol (orange), or ethanol (dark blue) as a carbon source (C, left). For each assay, a mutant *TDH3* genotype (yellow) was mixed with the common competitor genotype (green) (C, middle). The relative frequency of YFP- and GFP-fluorescent cells was recorded at regular intervals (∼6-8 generations) for ∼22 generations during log-phase growth (C, right). The fitness of the mutant genotype was estimated as the exponential of the slope of the natural log of the relative frequency of YFP and GFP cells over time. Relative fitness of the mutant strain was calculated by comparing the fitness of each mutant strain with the strain containing the wild-type *TDH3* promoter. D) Environment-dependent relationships between *TDH3* expression and fitness from Siddiq *et al*. [25] are shown for cells grown in media containing glucose (light blue), galactose (green), glycerol (orange), or ethanol (dark blue). Shading around each line indicates the 95% confidence interval. E-H) The relationship between median expression level (measured as a percentage of the median expression level driven by the wild-type *P*_*TDH3*_ allele) and expression noise (measured as a percentage of the standard deviation of expression driven by the wild-type *TDH3* allele) is shown for cells growing in glucose (E), galactose (F), glycerol (G) or ethanol (H). Within each environment, each of the 47 mutant *P*_*TDH3*_ alleles was designated as either “high noise” (orange) or “low noise” (green) based on whether its expression noise was greater than or less than, respectively, the expression noise expected for a genotype with that level of median expression based on a LOESS regression curve (black) using a value of 2/3 for the smoothing parameter. For each genotype in each environment, the difference between the observed expression noise and the expression noise predicted by the LOESS curve (i.e., residual of the LOESS regression, ‘Delta Noise’) provides a measure of expression noise independent of a genotype’s median expression level. I-L) The relationship between median expression level and fitness is shown separately for strains with low noise (green, Delta Noise < −1%) and high noise (orange, Delta Noise > +1%), for cells grown in media containing glucose (I), galactose (J), glycerol (K), or ethanol (L). In each case, the high- and low-noise LOESS regressions were estimated using a smoothing parameter *α* equal to 2/3. Shading around each line indicates the 95% confidence interval.

Because most mutations that change expression noise also change the average expression level [9,11,21], studying the fitness effects of changing expression noise requires controlling for the fitness effects of changing average expression level. Duveau *et al* [24] used the same data analyzed here for cells grown in glucose-based media to disentangle the effects of average expression level and noise on fitness by focusing on the changes in expression noise that were greater or less than expected based on the change in average expression level. We used the same approach here to compare the fitness effects of expression noise among environments. Briefly, in a given environment, we fitted a local regression model (LOESS) to infer the relationship between expression level and expression noise and used the model residuals to estimate the component of expression noise independent of mean expression level (Figure 2E-H). Using this LOESS regression, a strain was classified as having “high noise” if it had a positive residual (“Delta noise” in Figure 2E), meaning that it had higher noise than expected for a genotype with its median expression level, or “low noise” if it had a negative residual, meaning that it had lower noise than expected for a genotype with its median expression level (Figure 2E). We classified the 47 strains as high noise or low noise in each environment using the data from cells grown in each type of media (Figure 2E-H). 36%, 34%, 34%, and 38% of strains were classified as having “high noise” in glucose-, galactose-, glycerol-, and ethanol-based media, respectively. We then compared the fitness functions inferred using LOESS regression in each environment for strains categorized as having high or low expression noise in that environment. Visual inspection of these plots (Figures 2I-L) suggests that there are indeed differences in the fitness effects of expression noise among environments.

For cells grown in glucose, Duveau *et al. [24]* observed that genotypes with higher noise than expected based on their average expression level tended to be beneficial at lower expression levels that had lower fitness and deleterious at higher expression levels that were closer to the fitness optimum. To more formally test this observation, in each environment, we followed the same approach as Duveau *et al. [24]*, calculating the residual fitness (“Delta Fitness” in Figure 3A) for each strain as the difference between the observed fitness and the fitness expected for a genotype with that median expression level based on the empirically determined fitness function (Figure 3A-D). We then reclassified the strains as either “close to the optimum” (if fitness was within 0.5% of the environment-specific optimum) or “far from the optimum” (if fitness was more than 0.5% from the environment-specific optimum) (Figure 3A-D). For each of these classes, a significant correlation between Delta Noise and Delta Fitness would suggest that mutations causing changes in expression noise affect fitness independent of the effects of changing the average expression level on fitness.

**Figure 3:**
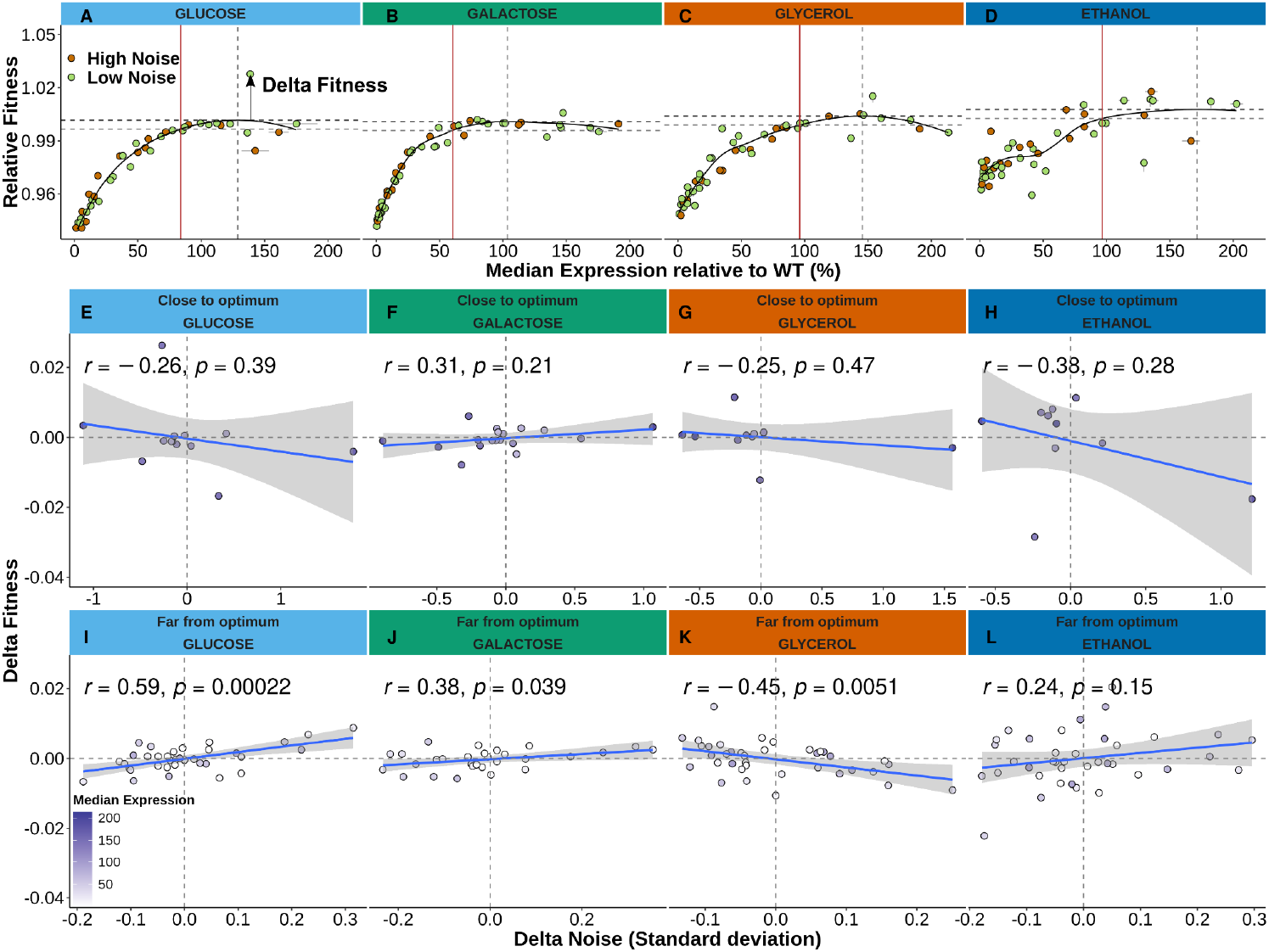
Expression noise can have different effects on fitness depending on median expression level. A-D) High-noise (green) and low-noise (orange) genotypes, defined as shown in Figure 2E-H, are overlaid with the fitness functions inferred for median expression levels of *TDH3* in glucose-(A, light blue), galactose-(B, green), glycerol-(C, orange), or ethanol-based (D, dark blue) media. In each environment, the expression optimum (vertical grey dotted line) indicates the *TDH3* expression level predicted to maximize fitness (dotted black horizontal line) from the LOESS regression of fitness on median expression (black curve). The expression level (vertical red line) at which the predicted fitness is 0.5% below the maximal fitness (dotted grey horizontal line) was chosen as the threshold separating genotypes with median activity ‘close to optimum’ (right-hand side of the red vertical line) from genotypes with median activity ‘far from optimum’ (left-hand side of the red vertical line). The residual of the LOESS regression (‘Delta Fitness’) is a measure of fitness effect independent of the median *TDH3* expression level. E-L) Relationship between Delta Noise and Delta Fitness for the *TDH3* mutant alleles for which median expression is close (E-H) or far from the optimum expression (I-L) in each environment [glucose (E, I), galactose (F, J), glycerol (G, K), and ethanol (H, L)]. Delta Noise, was estimated as the residual from the LOESS regression fitted to expression noise and median expression level. Delta Fitness was estimated as the residual from the LOESS regression fitted to relative fitness and median expression level. In each case (i.e., genotypes close to or far from the optimum in each environment), the best-fit linear model is shown in blue with grey shaded areas indicating 95% confidence intervals. The Pearson’s correlation coefficient and its p-value are shown in the upper left corner of each plot. In E-L, points are shaded based on the level of relative median *TDH3* expression.

For strains classified as close to the optimum, we saw no significant relationship between Delta Noise and Delta Fitness in any of the four environments (p > 0.05 in all cases, Figures 3E-H), which is consistent with the finding for glucose in Duveau *et al*. [30]. For strains far from the optimum, we observed that cells grown in glucose-based media had a statistically significant relationship (*r* = 0.59, p = 0.00022, Figure 3I), as previously reported in Duveau *et al*. [30], indicating that increasing expression noise is beneficial for genotypes with suboptimal expression levels. In galactose, we also observed a significant positive correlation (*r* = 0.38, p = 0.039, Figure 3J), suggesting the same relationship seen in glucose. (This pattern has also been reported for two other genes in *S. cerevisiae* [31].) In glycerol, we saw a significant negative correlation (*r* = -0.45, p = 0.0051, Figure 3K), suggesting that mutations decreasing (rather than increasing) expression noise are beneficial for genotypes with suboptimal expression levels. In ethanol, no significant correlation was observed (*r* = 0.24, p = 0.15, Figure 3L), suggesting no consistent fitness effects of increasing or decreasing expression noise in this environment.

Repeating these analyses using the variance divided by the median (similar to Fano factor) rather than the standard deviation to measure expression noise showed statistical support for the same relationships (Supplementary Figure 3). However, when the standard deviation scaled by the median (similar to CV) was used to measure expression noise, the only statistically significant relationship seen was for the strains far from the fitness optimum in glucose (Supplementary Figure 4).

### Simulating the fitness effects of expression noise

To investigate factors that might explain differences in the relationship between fitness and expression noise among environments, we used simulations of single-cell populations similar to those used in Duveau *et al. [24]*. These simulations compared the relative frequency of two genotypes during 600 minutes of simulated growth, providing a measure of their relative fitness. Each genotype was assumed to have its own normal distribution of expression levels among cells with that genotype. For the reference genotype included in each simulation, expression levels of single cells were drawn from a normal distribution. Expression levels of single cells for other genotypes were drawn from normal distributions with mean expression ranging from 0 to 2 and a standard deviation (i.e., expression noise) ranging from 0.05 to 0.8 (Figure 4A). A fitness function, describing the relationship (*f*(E)) between the gene expression level of an individual cell (E) and its doubling time (DT), was used for each simulation (Figure 4B). We first used a symmetric Gaussian fitness function (Supplementary Table 11), which has a single optimal expression level, similar to what was used in Duveau et al. [26]. Fitness of a genotype was defined as 1/DT (i.e., cells dividing faster have higher fitness). During this time-based simulation, each initial mother cell divided after a period of time (step size = 200 minutes), producing two daughter cells (Figure 4C). A new expression level was then determined for each daughter cell by drawing randomly from the distribution of single-cell expression levels for that genotype (Figure 4A). After division, we selected a total of 20,000 cells for reproduction based on probabilities proportional to the fitness of each tested genotype (1/DT). This process was repeated for two more rounds until the end of the simulated time. The relative fitness of each genotype was estimated as the exponential of the slope of the natural logarithm of its frequency relative to the reference genotype over time (Figure 4D).

**Figure 4:**
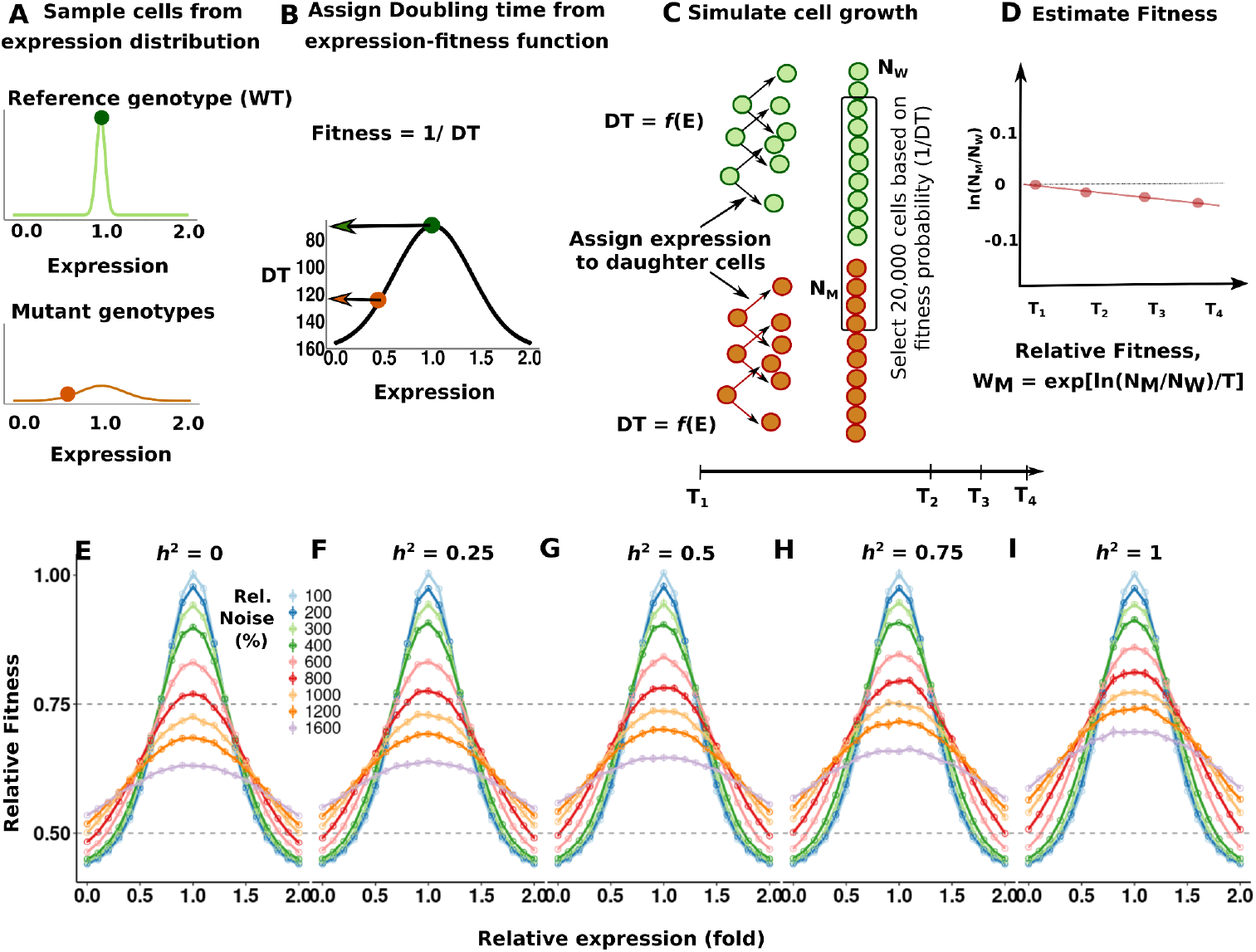
Heritability of expression between mother and daughter cells alters fitness effects of expression noise in simulations. A-D) Schematic representation of the population model simulating competitive growth for single gene expression noise. Each simulation was run for a short fixed duration (600 minutes) where variability in expression level (expression noise) impacting the single-cell division rate was the only determinant of population growth rate (fitness). In this model, the expression (E) of a gene was sampled for each genotype from a normal distribution ∼*N*(µ, σ^2^) where µ is the mean expression and σ is the standard deviation of expression (expression noise). A) The distributions of expression levels for cells with the wild-type (WT) reference genotype (green) and a mutant genotype (orange) are shown. In all simulations, the mutant genotype had a mean expression level between 0 and 2-fold of the mean expression level of the reference genotype (1.0), and the expression noise (standard deviation) ranged from 100% to 1600% of the expression noise of the reference genotype (0.05). B) The doubling time (DT) of a population was defined as a Gaussian function of expression: DT = *f*(E), with fitness represented by 1/DT. For each cell simulated from the reference (green dot) or mutant (orange dot) genotype, its doubling time was determined using this function. C) Each population simulation was initiated with 10,000 cells of the reference genotype (N_W_, green circles) and 10,000 cells of the mutant genotype (N_M_, orange circles), with the expression level of each cell determined by randomly drawing from that genotype’s distribution of expression levels shown in panel A. At each discrete time step (step size = 200 minutes), 20,000 cells were selected for reproduction based on a probability proportional to their fitness (1/DT). Each mother cell selected then produced two daughter cells. The expression level of each daughter cell was assigned as a weighted mix of the mother cell’s expression level, multiplied by its heritability, and an expression level randomly drawn from the distribution of expression levels for that genotype. Note that in simulations where the heritability was 0, expression of each daughter cell was determined solely by a random draw from the distribution of expression levels for that genotype. D) At the end of each simulation, the relative fitness of the mutant genotype was determined by the exponential (exp) of the slope of the natural logarithm (ln) of the ratio of the total number of cells from the mutant (N_M_) and wild-type reference (N_W_) genotypes over time (T). E-I) The relative fitness of each mutant genotype (Y-axis) is shown for mutants with relative expression levels ranging from 0 to 2-fold of the wild-type reference genotype (X-axis) and relative expression noise (Rel. Noise) ranging from 100% to 1600% of the expression noise of the wild-type reference genotype (different colored lines, with legend in figure). Results are shown for population simulations using a Gaussian fitness function relating single-cell expression levels to doubling times with heritability between mother and daughter cells (*h*^*2*^) of 0 (E), 0.25 (F), 0.5 (G), 0.75 (H), or 1 (I). For each data point in each panel, the error bars show 95% confidence intervals for relative fitness, estimated from 5 replicate simulations. Note that when expression was highly heritable (*h*^*2*^ = 1, panel I), there was a significant fitness cost of increasing expression noise in genotypes with average expression levels close to the expression optimum (Welch two-sample t-test p-value = 1.69 × 10^-12^) whereas there was a significant fitness benefit of increasing expression noise when the average expression level was far from the optimum (Welch two-sample t-test p-value = 7.93 × 10^-8^ and 1.38 × 10^-10^ for above and below the optimum, respectively). Additional statistical tests comparing the simulated fitness landscapes shown in E-I are provided in Supplementary Table 12.

Running this simulation with genotypes having a mean expression level ranging from 0 to 2 and values of expression noise ranging from 50% to 800% of that for the reference strain showed that increasing expression noise was deleterious for genotypes with a mean expression level close to the fitness optimum but beneficial for genotypes with a mean expression level far from the optimum (above or below 50% of the optimum expression) (Figure 4E; see Supplementary Table 12 for the detailed test statistics). This matches the pattern of fitness effects for expression noise reported previously for cells grown in glucose [24] and reported here for the first time for cells grown in galactose. It also aligns with findings from analytical models [32]. However, it differs from the relationship between expression noise and fitness seen for cells grown in glycerol- and ethanol-based environments. Why do we see these differences in the fitness effects of expression noise in different environments?

### Heritability of expression levels between mother and daughter cells alters the impact of expression noise on fitness

As described above, the doubling time is longer for cells growing in glycerol- or ethanol-based media than for cells growing in glucose- or galactose-based media; however, changing the doubling time parameter cannot explain the differences in fitness effects of expression noise across different mean expression levels in our simulations because doubling time acts simply as a scalar in this model. Differences in doubling time might, however, affect the similarity of expression levels *in vivo* between mother and daughter cells, which we consider here as heritability (*h*^*2*^). When assuming no heritability of expression level between mother and daughter cells (*h*^*2*^ = 0), the expression level of the daughter cell was simulated by a random draw from the expression distribution of that genotype, as was done in our initial simulation and in Duveau *et al*. [24]. To simulate the expression level of the daughter cell, assuming that it is completely inherited from the mother cell (*h*^*2*^ = 1), the expression level of the daughter cell was held equal to the expression level of the mother cell throughout the simulation. In such a case, with the same symmetric Gaussian fitness function, the fitness cost of increasing expression noise in genotypes with average expression levels close to the expression optimum was reduced (Figure 4I). For genotypes with an average expression level far from the optimum, increasing expression noise became even more beneficial than in simulations assuming no heritability (Figure 4I).

We also considered intermediate levels of heritability (i.e., *h*^*2*^ = 0.25, 0.5, or 0.75), which we think are likely the most realistic, given that *S. cerevisiae* cells divide by budding. To do so, we ran simulations where the expression level of each daughter cell was a weighted mixture of the mother cell’s expression and the expression level drawn randomly from that genotype’s distribution of expression levels. In these cases, the fitness costs and benefits for cells with average expression close to and far from the expression optimum, respectively, were intermediate and scaled with heritability (Figures 4F-H). In no case, however, did adding heritability into our simulations alter the relationship between expression noise and fitness in a way that was similar to the differences we saw between cells growing in fermentable (glucose, galactose) and non-fermentable (glycerol, ethanol) environments. For example, none of these simulations produced a pattern where increasing expression noise was deleterious for genotypes with average expression levels far from the optimum, as was seen for cells grown in glycerol-based media (Figure 3K).

### The shape of the relationship between gene expression level and fitness alters the fitness effects of mutations that modulate expression noise

With differences in doubling times and potential differences in heritability unable to explain the differences in fitness effects of expression noise that we observed among environments, we next considered the effect of differences in the shape of fitness functions, which reflects the sensitivity of fitness to changes in expression level. To do so, we performed simulations in which we replaced the symmetric, Gaussian fitness function (Figure 5A) with an asymmetric relationship between mean expression level and doubling time. Specifically, we used a lognormal fitness function (Figure 5B) and simulated populations assuming no heritability in expression level between mother and daughter cells. This lognormal function had a single optimal expression level, similar to the Gaussian function (Supplementary Table 11). We found that the fitness effects of increasing expression noise were again conditioned on the average expression level of a genotype, with increased expression noise associated with increased fitness when genotypes had a mean expression level that was far from the optimum (Figure 5E). However, the average expression level at which increasing expression noise transitioned from being beneficial to deleterious occurred at different average expression levels and distances from the fitness optimum with the Gaussian and lognormal fitness functions (Figures 5E and F). The point of this transition also differed as expression increased or decreased from the optimum with the asymmetric lognormal fitness function (Figure 5F). Interestingly, with this lognormal fitness function, the average expression level conferring the highest fitness among genotypes (i.e., the optimal expression) differed for genotypes with different levels of expression noise (Figure 5F).

**Figure 5:**
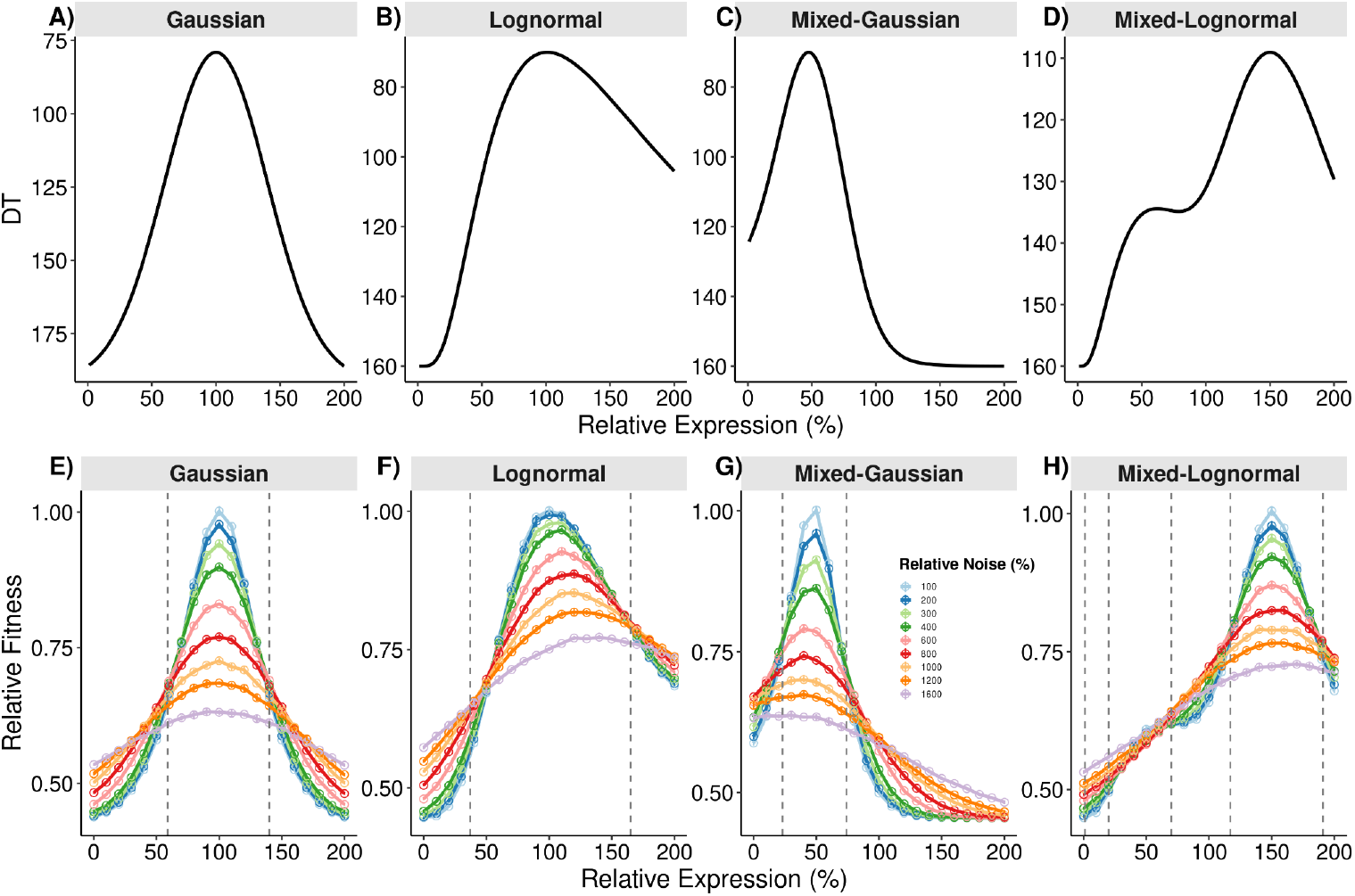
Population simulations show that different shapes of fitness functions change the effects of expression noise on fitness. A-D) DT = *f*(Expression) represents how expression of a single gene in a single cell relates to its doubling time (DT). Expression levels were defined as percentages relative to the mean expression level of a reference strain. This relationship was defined as either a Gaussian (A) or Lognormal (B) function or a mixture of two different Gaussian (C) or Lognormal functions (D). E-H) The relative fitness of each mutant genotype (Y-axis) is shown for mutants with relative expression levels ranging from 0 to 200% of the reference genotype (X-axis); relative expression noise ranging from 100% to 1600% of the expression noise of the wild-type reference genotype (different colored lines, with legend in figure); and the relationship between expression level and doubling time defined using a Gaussian (E), Lognormal (F), Mixed-Gaussian (G), or Mixed-Lognormal (H) function. For each data point in each panel, the error bars show 95% confidence intervals for relative fitness, estimated from 5 replicate simulations. For each fitness function (A-D), inflection points (i.e., where the concavity of the curve changes) were determined by the second derivative of the function using the smoothing spline method and are marked by grey dotted vertical lines in the corresponding simulation plots (E-H).

To determine how more complex fitness functions alter the relationship between expression noise and fitness, we also performed simulations in which the fitness functions were either a mixture of two Gaussian or two lognormal functions (Supplementary Table 11). Both of these fitness functions were asymmetrical, and the mixture of two lognormal functions had a sub-optimal fitness peak in addition to the overall fitness optimum (Figures 5C and D, respectively). Once again, we observed that low expression noise conferred the highest fitness for genotypes with average expression levels close to the fitness optimum and that increases in expression noise were beneficial among genotypes with average expression levels farthest from the optimum (Figures 5G and H). However, we also noticed in the simulation using a mixed Gaussian function (Figure 5G) that when the average expression level was below the optimum, genotypes with intermediate levels of noise (800% relative to a genotype with a standard deviation of 0.05) had higher fitness than genotypes with either lower (100%) or higher (1600%) expression noise. The simulation using mixed lognormal functions also showed that genotypes with intermediate (800%) noise had the highest fitness for some average expression levels, but in this case, it occurred when the average expression level was above (rather than below) the optimum as well as when it was in the valley between the two fitness peaks (Figure 5H). These regions of the fitness function represent average expression levels for which fitness was maximized by a specific level of expression noise. That is, for genotypes with this average expression level, the optimal noise level effectively ‘bridges the gap’ between a suboptimal average expression level and the fitness peak. Any deviation from this balance is deleterious: genotypes with lower noise fail to produce enough cells with expression close to the fitness optimum, whereas genotypes with higher noise produce too many cells with expression conferring lower fitness.

Considering the fitness landscapes simulated with all four types of fitness functions (Gaussian, lognormal, mixed-Gaussian, and mixed-lognormal) together, we noted that genotypes near the optimum with higher expression noise always showed a lower maximal fitness (Figures 5E-H), presumably because noisier genotypes have more individual cells with suboptimal expression levels. The fitness functions for genotypes with high levels of expression noise were also always flatter (i.e., spanned a narrower range of fitness values) than genotypes with lower levels of expression noise (Figures 5E-H), consistent with selection being less effective for changing the average expression level when expression is noisy [33]. Comparing the fitness effects of changing expression noise to the shape of the fitness function, we found that mutations increasing expression noise are generally predicted to be beneficial for genotypes with an average expression level that corresponds to an area of the fitness function that is convex (curving upward or leveling off, as seen near the tails of a Gaussian distribution where expression is far from optimum). Mutations increasing expression noise are generally predicted to be deleterious for genotypes with an average expression level that corresponds to an area of the fitness function that is concave (i.e., curving downward, as typically seen near a fitness peak). These observations are consistent with the mathematical property of nonlinear functions known as Jensen’s inequality [34], a connection that has also recently been made in a theoretical study of expression noise by Weinreich et al. [29]. Overall, the extent of expression noise, the shape of the fitness function, and the distance between a genotype’s average expression level and the optimal expression level work together to determine the fitness effects of expression noise, as seen in the simulations using mixtures of Gaussian or lognormal functions (Figures 5G and H).

Regardless of the shape of the fitness function used, including heritability of expression levels between mother and daughter cells into each of these simulations caused smaller differences in fitness among genotypes with differences in expression noise when the average expression level was close to the optimum and larger differences in fitness among genotypes with differences in expression noise when the average expression level was far from the optimum (Supplementary Figure 5). This observation suggests that selection favoring higher expression noise in genotypes where the average expression level is far from the optimum will be most effective when heritability of expression between mother and daughter cells is high.

### Conclusions

We found that expression noise driven by the *S. cerevisiae TDH3* promoter was plastic (i.e., differed for the same promoter allele among environments) and tended to be lower in environments in which cells divide faster. We also found that the fitness effects of expression noise seen previously for cells growing in glucose—namely, that increasing expression noise is deleterious for genotypes with average expression levels close to the fitness optimum but beneficial for genotypes with average expression levels far from the optimum [26]—were also seen for cells growing in galactose but not for cells growing in glycerol or ethanol. Simulations of single cells suggested that differences in the shape of the function relating expression level to fitness are most likely to explain these observations. Specifically, we found that increasing expression noise was beneficial when the average expression level was in a convex region of the fitness function and deleterious when the average expression level was in a concave region of the fitness function, consistent with Jensen’s inequality from mathematics. We expect that these observations will also extend to expression of genes other than *TDH3*, as well as to traits other than gene expression, because our model did not assume any properties specific to *TDH3*. Incorporating heritability of expression levels from mother to daughter cells in our simulations suggested that higher heritability increases the benefits of increasing expression noise in convex regions of the fitness function and decreases the fitness cost of higher noise in concave regions. Taken together, these findings show how changes in the environment can shape the evolution of gene expression by altering not only mean expression levels and expression noise but also the selective consequences of expression noise, which result from environment-specific relationships between expression level and fitness.

## Materials and Methods

### Yeast strains

This study uses measures of expression level, expression noise, and fitness for various alleles of the *TDH3* promoter (*P*_*TDH3*_) from Siddiq *et al*. [25]. These data come from analysis of a haploid (mating-type “a”) strain containing either a mutant or reference allele of the *P*_*TDH3*_*-YFP* construct at the *HO* locus and a wild type P_*TDH3*_ at the native locus (progenitor strain YPW1002). Each of these mutant strains carried a mutation(s) in the *TDH3* promoter driving YFP expression at the *HO* locus. To assess the effects of overexpression, eight of the strains contained two copies of *P*_*TDH3*_*-YFP* at the *HO* locus, with a *URA3* gene separating the two copies. Relative to the BY strain of *S. cerevisiae*, all of these strains also had an A-293-G mutation in the *TDH3* promoter, which has a negligible effect on *TDH3* expression [24]. To account for autofluorescence, strain YPW978 (restocked as strain YPW3858), which does not contain a YFP reporter construct, was also analyzed.

Another set of yeast strains, made in the haploid genetic background of strain YPW1001 (mating type “α”), in which the endogenous *TDH3* promoter carried the same mutation(s) as the mutant *P*_*TDH3*_*-YFP* strains, was used to measure the fitness effects of changing *TDH3* expression. These strains also had an unmutated *P*_*TDH3*_*-YFP* construct at the *HO* locus that allowed them to be distinguished from the reference strain carrying a *P*_*TDH3*_*-GFP* construct at the *HO* locus (strain YPW1160) during the competitive growth assays used to estimate fitness. There was no significant difference in fluorescence between strains YPW1002 and YPW1001, suggesting that the few genetic differences between the genetic backgrounds used to measure expression and fitness (e.g., differences in mating type) did not significantly affect *TDH3* promoter activity [24].

### Environmental conditions

Four types of growth media that differed only in terms of their carbon source were used to generate the data analyzed in this study [25]: rich medium (10 g/l yeast extract and 20 g/l peptone dissolved in water) complemented with glucose (YPD, 20g/l dextrose monohydrate), galactose (YPGal, 20 g/l galactose), glycerol (YPG, 30 ml of 99% glycerol per litre), and ethanol (YPE, 50 ml of 99% ethanol per liter). Media were filter-sterilized before experiments.

### Measuring YFP fluorescence using flow cytometry

All strains were initially revived from glycerol stocks by growing them on a YPG plate (10 g/l yeast extract, 20 g/l peptone, 30 ml of 99% glycerol per liter, 20 g/l agar) for two days. Later, the set of 51 strains (47 strains carrying mutant alleles of the *TDH3* promoter and 4 control strains) was randomly arrayed into a 96-well plate containing YPD media. Cells were maintained in suspension in plates by growing in a shaking incubator for 22 hours at 30 °C in YPD media. Cells were then diluted in one of the four types of media (glucose, galactose, glycerol, or ethanol) and acclimatized by diluting to fresh media every 12 hours for 36 hours. Cell density was measured before each dilution, and a dilution factor was estimated in such a way that at the end of 12 hours of growth, the cell density was ∼5 × 10^-6^ cells/ml. These conditions maintained the cells in exponential growth. After 36 hours, the samples were diluted to 2.5 × 10^-6^ cells/ml with Phosphate Buffer (PBS), and fluorescence (as well as an estimate of cell size) was captured for 20,000 events per well on a BD Accuri C6 flow cytometer coupled to a HyperCyt autosampler (IntelliCyt Corp.). YFP fluorescence was measured using a 530/30 optical filter on this flow cytometer.

### Analysis of flow cytometry data

Flow cytometry data were analyzed in the R environment using the flowCore [35] and flowClust [36] packages, as described by Duveau *et al*. [24]. Briefly, single events were isolated using automated gating for forward scatter height (FSC-H) and area (FSC-A). The intensity of the fluorescence signal was scaled by cell size estimated using these parameters, and the YFP signal was adjusted for autofluorescence using the signal from the YPW978 strain lacking a YFP reporter construct. The median YFP expression level for each genotype in each replicate was estimated as the residual of the linear model fitting for positional effect using the YPW1002 reference strain as a control strain. Finally, the relative YFP expression for each genotype was calculated after scaling the median YFP expression level by the corresponding mean of the median YFP expression for the relevant unmutated reference strain (YPW1002 for strains with a single copy of the *TDH3* promoter and YPW2675 for strains with two copies of the *TDH3* promoter).

### Measuring fitness

To measure the relative fitness of different *TDH3* promoter alleles in different environments, Duveau *et al*. [24] and Siddiq *et al*. [25] used head-to-head competition assays between each strain with mutations in the native *TDH3* promoter (marked with *P*_*TDH3*_*-YFP* at the *HO* locus) and a reference strain with an unmutated *TDH3* promoter (marked with *P*_*TDH3*_*-GFP* at the *HO* locus, strain YPW1160) in each of the four environments. Relative growth during log-phase was used as a proxy for relative fitness.

For each competitive growth experiment, YFP and GFP strains were thawed from glycerol stocks, grown on YPG agar media, and then transferred to liquid YPD media. Cells were grown at 30°C for 24 h while shaking. Equal volumes of YFP and GFP cell cultures were added to wells of four 96-well plates (one containing media with each carbon source), and samples were diluted to achieve a density of ∼5 × 10^-6^ cells/ml after 12 h of growth. After three rounds of cell density measurement, dilution in fresh media, and 12 hours of growth to acclimatize in the specific growth medium, the competitive growth assay was started, repeating three additional cycles of dilution and growth. The ratio of YFP- to GFP-expressing cells in each well was recorded before each of these final three rounds of dilution as well as at the end of the experiment using flow cytometry. Approximately 75,000 events were recorded for each sample on a BD Accuri C6 flow cytometer, using a 488-nm laser for excitation and two different optical filters (510/10 and 585/40) to acquire fluorescence.

These data were used to determine the number of cell generations that occurred during the three dilution cycles, using the median number of generations for all samples grown on the same 96-well plate. Next, the relative fitness of the YFP strain was estimated as the exponential of the slope of the natural logarithm of the ratio of YFP-positive over GFP-positive cells by fitting a linear regression model with the number of generations across the four time points. For each mutant, the fitness relative to the GFP strain was then divided by the mean fitness for all replicates of the reference strain YPW1189 (for single-copy *P*_*TDH3*_ variants) or YPW2682 (for double-copy *P*_*TDH3*_ variants). Finally, we estimated the mean relative fitness and standard deviation of each mutant strain across replicates. The same protocol was used for all environments except that the amount of time cells were allowed to grow before each dilution was adjusted to account for environment-specific doubling times.

### Simulations of single-cell populations

We performed stochastic simulations of the growth of clonal cell populations spanning a wide range of mean expression levels (0% to 200% relative to wild-type) and expression noise values (5% to 800% relative to wild-type noise) of a single gene. We simulated the evolution of gene expression in two competing genotypes (A and B) using an individual-based model implemented in Python. All simulations were run using the DEAP package (version 1.4.3) in the Python (version 3.13.9) environment as a short fixed duration (600 minutes with a step size of 200 minutes) experiment where variability in expression level impacting the single cell division rate was the only determinant of population growth rate. The behavior of each genotype was determined by a normal distribution *N* of expression levels for the focal genotype described by its mean (µ) and variance (σ^2^) and a fitness function, Doubling Time (DT) = *f*(E). We tested several different fitness functions: (1) Gaussian function (symmetric), (2) lognormal function (asymmetric), (3) mixed distribution of two different Gaussian functions (asymmetric), or (4) mixed distribution of two different lognormal functions (asymmetric, with more than one local fitness optimum).

In each run of the simulation, 10,000 cells of the reference (wild-type) genotype were generated, with expression in each cell determined by a random draw from a normal distribution (mean = fitness optimum; standard deviation = 0.05). The DT of each cell was then estimated based on its expression level using the fitness function. For the competing genotype (mutant genotype), we again generated 10,000 cells with expression of each cell drawn from a normal distribution with a mean expression level ranging from 0 to 2 and a standard deviation ranging from 0.05 to 0.8. At each discrete time step (200 minutes), individuals were selected for reproduction based on a probability proportional to their fitness, and each mother cell selected produced two daughter cells. We first simulated genotypes where expression heritability was zero by randomly assigning the expression of daughter cells from the same distribution of expression levels sampled in the initial population. The relative fitness of each genotype was estimated by comparing the total number of mutant cells (N_M_) and the total number of wild-type cells (N_W_) obtained from simulating the growth for 600 minutes, which is 3 time intervals (T), using the following equation:

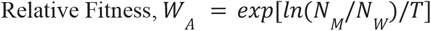

Next, to understand the impact of heritability in expression level between mother and daughter cells on the relationship between gene expression noise and fitness, we tested different levels of expression heritability (*h*^2^ ranging from 0 to 1) while assigning the expression of daughter cells using the following equation and then estimated the relative fitness as described above:

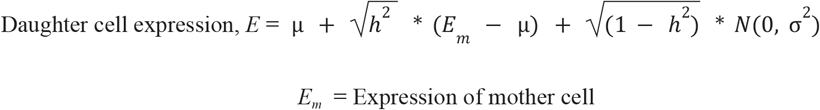

## Data accessibility

Flow Cytometry data (.fcs files) used in this study can be downloaded from http://flowrepository.org/ with Repository ID FR-FCM-Z8EK for the expression data and Repository ID FR-FCM-Z8EQ for the data used to measure fitness from competitive growth assays. Code used to analyze data, prepare figures, and perform simulations can be found at https://github.com/tahia/Noise_Plasticity.

## Acknowledgements

We thank the members of the Wittkopp Lab, including Erick Bayala Rodriguez, Rebecca McAvoy, and Ayushi Dasgupta, for helpful discussion and comments on this work. We also thank the University of Michigan Flow Cytometry Core facility for access to equipment and advice and the University of Michigan Advanced Research Computing facility for access to the Great Lakes computing cluster.

## Author’s Contributions

F.D., M.A.S., T.H., and P.J.W. planned and designed the research. F.D. collected the data. T.H. and M.A.S. analyzed the data. T.H. and P.J.W. wrote the manuscript, with review and contributions from all the authors.

## References

1. Scheiner SM. 1993 Genetics and evolution of phenotypic plasticity. Annu. Rev. Ecol. Syst. 24, 35–68.

2. Viney M, Reece SE. 2013 Adaptive noise. Proc. Biol. Sci. 280, 20131104.

3. Elowitz MB, Levine AJ, Siggia ED, Swain PS. 2002 Stochastic gene expression in a single cell. Science 297, 1183–1186.

4. Fraser HB, Hirsh AE, Giaever G, Kumm J, Eisen MB. 2004 Noise minimization in eukaryotic gene expression. PLoS Biol. 2, e137.

5. Metzger BPH, Yuan DC, Gruber JD, Duveau F, Wittkopp PJ. 2015 Selection on noise constrains variation in a eukaryotic promoter. Nature 521, 344–347.

6. Des Marais DL, Hernandez KM, Juenger TE. 2013 Genotype-by-environment interaction and plasticity: Exploring genomic responses of plants to the abiotic environment. Annu. Rev. Ecol. Evol. Syst. 44, 5–29.

7. Smith EN, Kruglyak L. 2008 Gene-environment interaction in yeast gene expression. PLoS Biol. 6, e83.

8. Raser JM, O’Shea EK. 2004 Control of stochasticity in eukaryotic gene expression. Science 304, 1811–1814.

9. Newman JRS, Ghaemmaghami S, Ihmels J, Breslow DK, Noble M, DeRisi JL, Weissman JS. 2006 Single-cell proteomic analysis of S. cerevisiae reveals the architecture of biological noise. Nature 441, 840–846.

10. Duveau F, Yuan DC, Metzger BPH, Hodgins-Davis A, Wittkopp PJ. 2017 Effects of mutation and selection on plasticity of a promoter activity in Saccharomyces cerevisiae. Proc. Natl. Acad. Sci. U. S. A. 114, E11218–E11227.

11. Hornung G, Bar-Ziv R, Rosin D, Tokuriki N, Tawfik DS, Oren M, Barkai N. 2012 Noise-mean relationship in mutated promoters. Genome Res. 22, 2409–2417.

12. Krieger G, Lupo O, Levy AA, Barkai N. 2020 Independent evolution of transcript abundance and gene regulatory dynamics. Genome Res. 30, 1000–1011.

13. Gasch AP, Spellman PT, Kao CM, Carmel-Harel O, Eisen MB, Storz G, Botstein D, Brown PO. 2000 Genomic expression programs in the response of yeast cells to environmental changes. Mol. Biol. Cell 11, 4241–4257.

14. Li XC, Fay JC. 2017 Cis-Regulatory Divergence in Gene Expression between Two Thermally Divergent Yeast Species. Genome Biol. Evol. 9, 1120–1129.

15. Kita R, Venkataram S, Zhou Y, Fraser HB. 2017 High-resolution mapping of cis-regulatory variation in budding yeast. Proc. Natl. Acad. Sci. U. S. A. 114, E10736–E10744.

16. Lehner B. 2010 Conflict between noise and plasticity in yeast. PLoS Genet. 6, e1001185.

17. Field Y, Kaplan N, Fondufe-Mittendorf Y, Moore IK, Sharon E, Lubling Y, Widom J, Segal E. 2008 Distinct modes of regulation by chromatin encoded through nucleosome positioning signals. PLoS Comput. Biol. 4, e1000216.

18. Tirosh I, Barkai N. 2008 Two strategies for gene regulation by promoter nucleosomes. Genome Res. 18, 1084–1091.

19. Zenklusen D, Larson DR, Singer RH. 2008 Single-RNA counting reveals alternative modes of gene expression in yeast. Nat. Struct. Mol. Biol. 15, 1263–1271.

20. Keren L et al. 2016 Massively parallel interrogation of the effects of gene expression levels on fitness. Cell 166, 1282–1294.e18.

21. Keren L, van Dijk D, Weingarten-Gabbay S, Davidi D, Jona G, Weinberger A, Milo R, Segal E. 2015 Noise in gene expression is coupled to growth rate. Genome Res. 25, 1893–1902.

22. Wittkopp PJ. 2023 Contributions of mutation and selection to regulatory variation: lessons from the Saccharomyces cerevisiae TDH3 gene. Philos. Trans. R. Soc. Lond. B Biol. Sci. 378, 20220057.

23. Metzger BPH, Duveau F, Yuan DC, Tryban S, Yang B, Wittkopp PJ. 2016 Contrasting Frequencies and Effects of cis- and trans-Regulatory Mutations Affecting Gene Expression. Mol. Biol. Evol. 33, 1131–1146.

24. Duveau F, Hodgins-Davis A, Metzger BP, Yang B, Tryban S, Walker EA, Lybrook T, Wittkopp PJ. 2018 Fitness effects of altering gene expression noise in Saccharomyces cerevisiae. Elife 7, e37272.

25. Siddiq MA, Duveau F, Wittkopp PJ. 2024 Plasticity and environment-specific relationships between gene expression and fitness in Saccharomyces cerevisiae. Nat. Ecol. Evol. 8, 2184–2194.

26. Duveau F, Toubiana W, Wittkopp PJ. 2017 Fitness Effects of Cis-Regulatory Variants in the Saccharomyces cerevisiae TDH3 Promoter. Mol. Biol. Evol. 34, 2908–2912.

27. Gancedo JM. 1998 Yeast carbon catabolite repression. Microbiol. Mol. Biol. Rev. 62, 334–361.

28. Liti G. 2015 The fascinating and secret wild life of the budding yeast S. cerevisiae. Elife 4. (doi:10.7554/eLife.05835)

29. Weinreich DM, Sgouros T, Raynes Y, Burtsev H, Chang E, Rajakumar S, Bravo IG, Petak C. 2026 Wanted: A population genetic theory of biological noise regulation. PLoS Genet. 22, e1012066.

30. Duveau F, Hodgins-Davis A, Metzger BP, Yang B, Tryban S, Walker EA, Lybrook T, Wittkopp PJ. 2018 Fitness effects of altering gene expression noise in Saccharomyces cerevisiae. Elife 7, e37272.

31. Schmiedel JM, Carey LB, Lehner B. 2019 Empirical mean-noise fitness landscapes reveal the fitness impact of gene expression noise. Nat. Commun. 10, 3180.

32. Tănase-Nicola S, ten Wolde PR. 2008 Regulatory control and the costs and benefits of biochemical noise. PLoS Comput. Biol. 4, e1000125.

33. Wang Z, Zhang J. 2011 Impact of gene expression noise on organismal fitness and the efficacy of natural selection. Proceedings of the National Academy of Sciences 108, E67–E76.

34. Ruel JJ, Ayres MP. 1999 Jensen’s inequality predicts effects of environmental variation. Trends Ecol. Evol. 14, 361–366.

35. Ellis B, Haaland P, Hahne F, Meur L, Gopalakrishnan N, Spidlen N, Jiang J, Finak M. 2026 flowCore: flowCore: Basic structures.

36. Lo K, Hahne F, Brinkman RR, Gottardo R. 2009 flowClust: a Bioconductor package for automated gating of flow cytometry data. BMC Bioinformatics 10, 145.

