## Supplementary Figures for "Environmental impacts on gene expression noise and its relationship with fitness"

**Supplementary Materials for**
### Environmental impacts on gene expression noise and its relationship with fitness

by Taslima Haque, Mohammad A. Siddiq, Fabien Duveau, and Patricia J. Wittkopp

**Supplementary Figures:**

Supplementary Figure 1. Plasticity of expression noise measured by Fano factor
Supplementary Figure 2. Plasticity of expression noise measured by CV
Supplementary Figure 3. Fitness effects of gene expression noise measured by Fano factor
Supplementary Figure 4. Fitness effects of gene expression noise measured by CV
Supplementary Figure 5: Impacts of heritability on the fitness effects of expression noise

**Supplementary Tables:** (provided as separate sheets in a single Excel file with a README tab)

Supplementary Table 1. Description of the yeast strains used in this study
Supplementary Table 2. Expression data for *TDH3* promoter mutant alleles grown in Glucose
Supplementary Table 3. Expression data for *TDH3* promoter mutant alleles grown in Galactose
Supplementary Table 4. Expression data for *TDH3* promoter mutant alleles grown in Glycerol
Supplementary Table 5. Expression data for *TDH3* promoter mutant alleles grown in Ethanol
Supplementary Table 6. Summary statistics of the expression of *TDH3* mutants across four environments
Supplementary Table 7. Test statistics of mutant-allele dependent noise variation
Supplementary Table 8. Test statistics for expression plasticity among mutant *TDH3* promoter alleles across four environments
Supplementary Table 9. Replicate-specific fitness measures from mutant *TDH3* promoter alleles in four environments
Supplementary Table 10. Fitness estimates of mutant *TDH3* promoter alleles in four environments
Supplementary Table 11. Single-cell population simulation parameters
Supplementary Table 12: Test statistics of simulated fitness variation

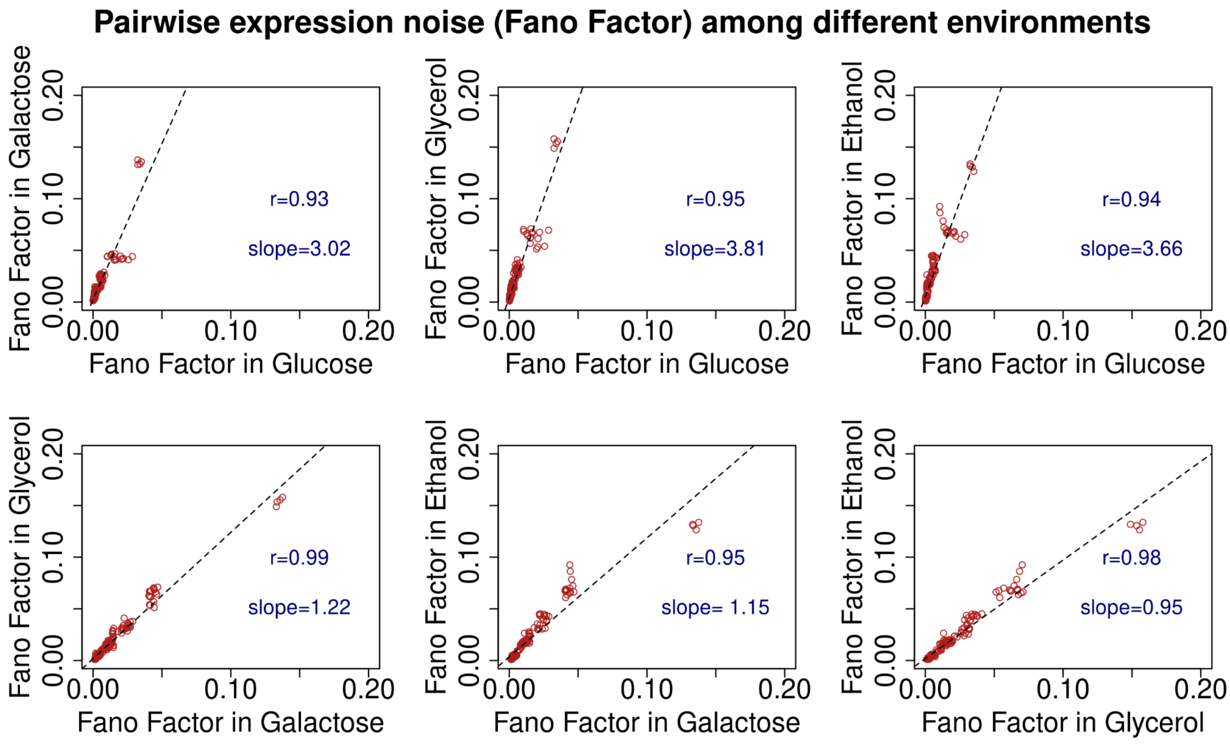


**Supplementary Figure 1. Plasticity of expression noise measured by Fano factor.** Fano Factor (squared standard deviation scaled by median expression level) of YFP expression is plotted as a measure of expression noise for the 47 $P_{TDH3}$-*YFP* reporter genes in all six pairs of environments. The X and Y axes indicate the specific environments (glucose-, galactose-, glycerol-, or ethanol-based media) compared in each plot. Dotted lines show the linear regression for each pair of environments, and the regression slope is indicated on each plot. The effects of *TDH3* promoter mutations on expression noise were highly correlated (Pearson's $r > 0.93$) between all six pairs of environment contrasts, as was also seen when standard deviation was used to measure expression noise (Figure 1E-J).

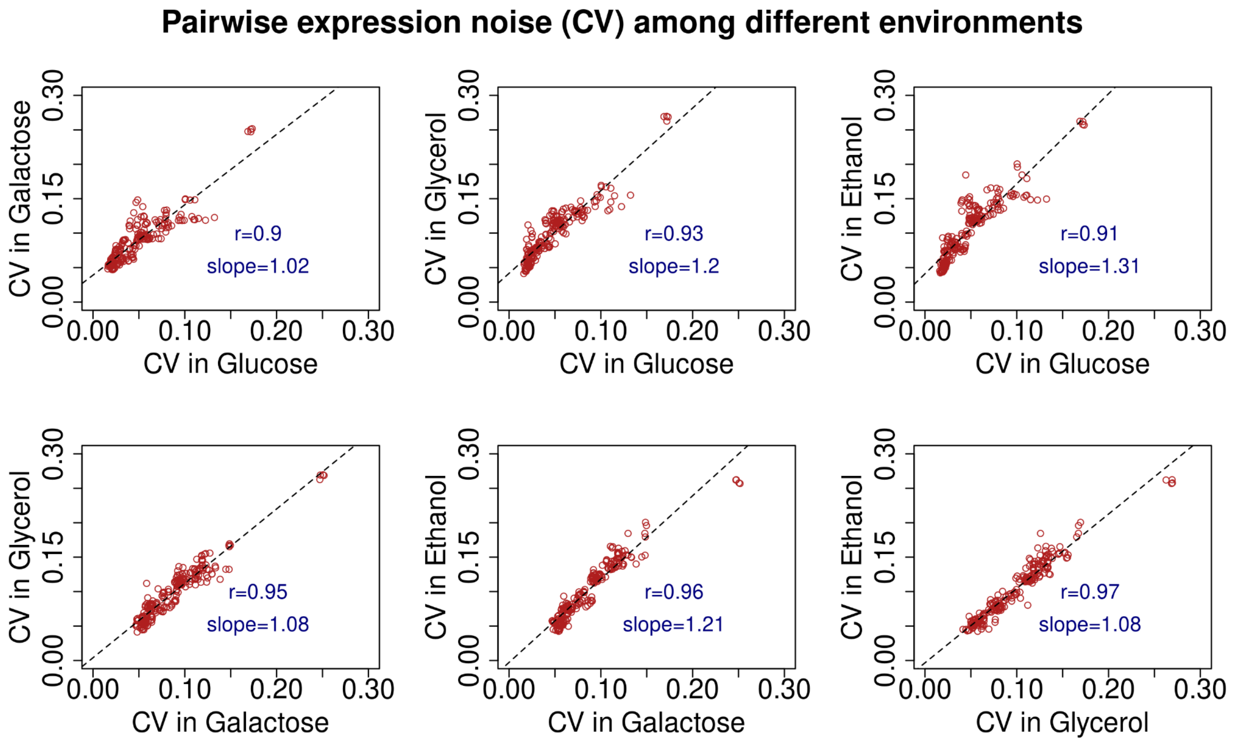


**Supplementary Figure 2. Plasticity of expression noise measured by CV.** Coefficient of variation (CV, standard deviation scaled by expression median) of YFP expression is plotted as a measure of expression noise for the 47 $P_{TDH3}$-*YFP* reporter genes in all six pairs of environments. The X and Y axes indicate the specific environments (glucose-, galactose-, glycerol-, or ethanol-based media) compared in each plot. Dotted lines show the linear regression for each pair of environments, and the regression slope is indicated on each plot. The effects of *TDH3* promoter mutations on expression noise were highly correlated (Pearson's $r > 0.89$) between all six pairs of environment contrasts, as was also seen when standard deviation was used to measure expression noise (Figure 1E-J).

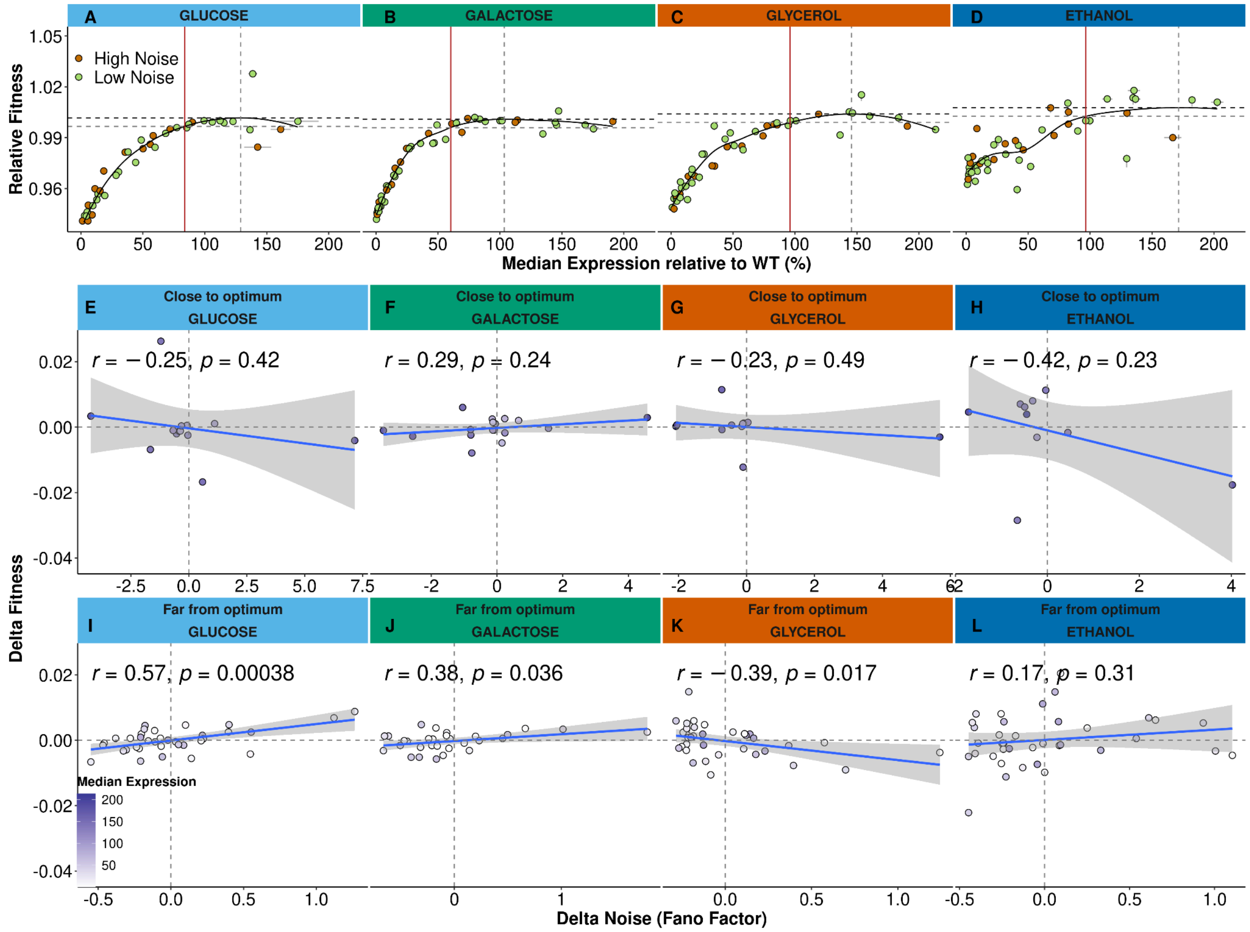


**Supplementary Figure 3**: **Fitness effects of gene expression noise measured by Fano Factor.** A-D) High-noise (green) and low-noise (orange) genotypes, defined as shown in Figure 2E-H, are overlaid with the fitness functions inferred for median expression levels of *TDH3* in glucose- (A, light blue), galactose- (B, green), glycerol- (C, orange), or ethanol-based (D, dark blue) media. In each environment, the expression optimum (vertical grey dotted line) indicates the *TDH3* expression level predicted to maximize fitness (dotted black horizontal line) from the LOESS regression of fitness on median expression (black curve). The expression level (vertical red line) at which the predicted fitness is 0.5% below the maximal fitness (dotted grey horizontal line) was chosen as the threshold separating genotypes with median activity 'close to optimum' (right-hand side of the red vertical line) from genotypes with median activity 'far from optimum' (left-hand side of the red vertical line). The residual of the LOESS regression ('Delta Fitness') is a measure of fitness effect independent of the median *TDH3* expression level. E-L) Relationship between Delta Noise and Delta Fitness for the *TDH3* mutant alleles for which median expression is close (E-H) or far from the optimum expression (I-L) in each environment [glucose (E, I), galactose (F, J), glycerol (G, K), and ethanol (H, L)]. Note that the Fano factor was used to measure expression noise here, whereas standard deviation was used to measure expression noise in Figure 2. Delta Noise, was estimated as the residual from the LOESS regression fitted to expression noise and median expression level. Delta Fitness was estimated as the residual from the LOESS regression fitted to relative fitness and median expression level. In each case (i.e., genotypes close to or far from the optimum in each environment), the best-fit linear model is shown in blue with grey shaded areas indicating 95% confidence intervals. The Pearson's correlation coefficient and its p-value are shown in the upper left corner of each plot. In E-L, points are shaded based on the level of relative median *TDH3* expression.

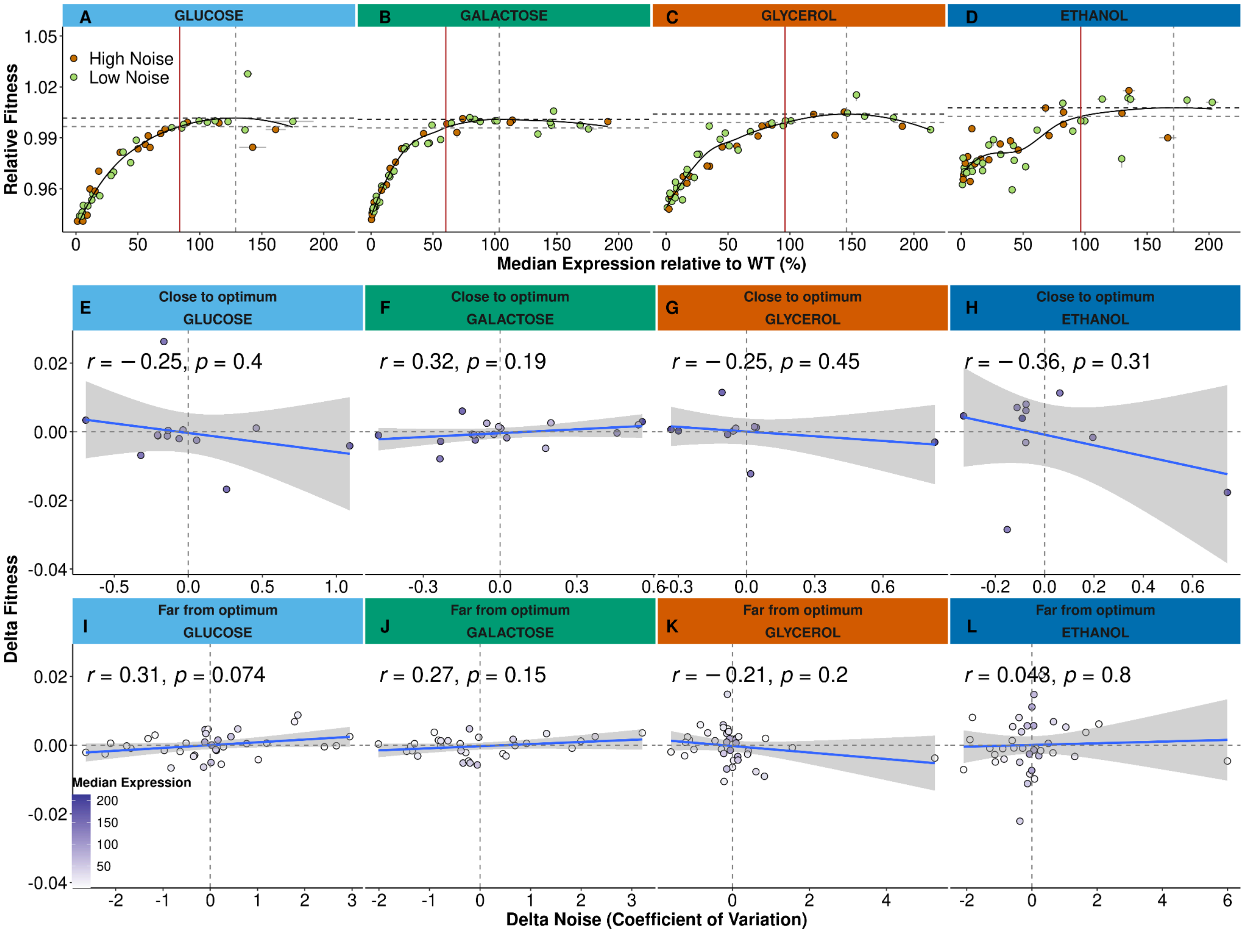


**Supplementary Figure 4**: **Fitness effects of gene expression noise measured by CV.** A-D) High-noise (green) and low-noise (orange) genotypes, defined as shown in Figure 2E-H, are overlaid with the fitness functions inferred for median expression levels of *TDH3* in glucose- (A, light blue), galactose- (B, green), glycerol- (C, orange), or ethanol-based (D, dark blue) media. In each environment, the expression optimum (vertical grey dotted line) indicates the *TDH3* expression level predicted to maximize fitness (dotted black horizontal line) from the LOESS regression of fitness on median expression (black curve). The expression level (vertical red line) at which the predicted fitness is 0.5% below the maximal fitness (dotted grey horizontal line) was chosen as the threshold separating genotypes with median activity 'close to optimum' (right-hand side of the red vertical line) from genotypes with median activity 'far from optimum' (left-hand side of the red vertical line). The residual of the LOESS regression ('Delta Fitness') is a measure of fitness effect independent of the median *TDH3* expression level. E-L) Relationship between Delta Noise and Delta Fitness for the *TDH3* mutant alleles for which median expression is close (E-H) or far from the optimum expression (I-L) in each environment [glucose (E, I), galactose (F, J), glycerol (G, K), and ethanol (H, L)]. Note that the coefficient of variation (CV)

factor was used to measure expression noise here, whereas standard deviation was used to measure expression noise in Figure 2. Delta Noise, was estimated as the residual from the LOESS regression fitted to expression noise and median expression level. Delta Fitness was estimated as the residual from the LOESS regression fitted to relative fitness and median expression level. In each case (i.e., genotypes close to or far from the optimum in each environment), the best-fit linear model is shown in blue with grey shaded areas indicating 95% confidence intervals. The Pearson's correlation coefficient and its p-value are shown in the upper left corner of each plot. In E-L, points are shaded based on the level of relative median *TDH3* expression.

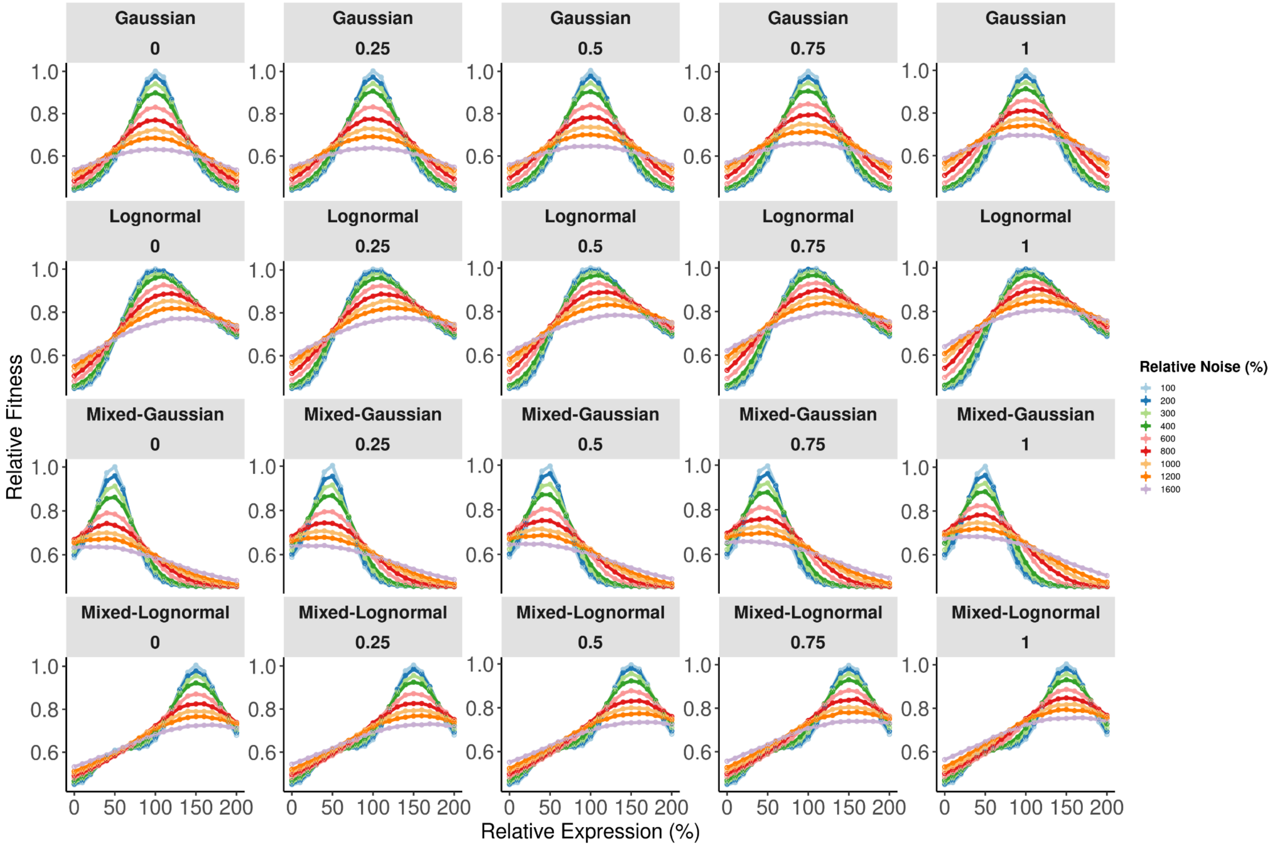


**Supplementary Figure 5**: **Impacts of heritability on the fitness effects of expression noise.** Relationships between relative fitness and mean expression level are shown from simulations of single-cell population growth of genotypes with different average expression levels, relative levels of noise, and heritability with four different fitness functions (Gaussian, Lognormal, Mixed-Gaussian, and Mixed-Lognormal). Relative expression levels ranged from 0 to 200% of wild-type. Relative noise ranged from 100% to 1600% of the reference standard deviation of 0.05. Heritabilities tested for each of the four models (shown under the model name in each case) were 0, 0.25, 0.5, 0.75, 1. Error bars indicate 95% confidence intervals for relative fitness estimated from 5 replicate simulations of each genotype under each set of conditions.
